# HIV-1 Vpu protein forms stable oligomers in aqueous solution via its transmembrane domain self-association

**DOI:** 10.1101/2023.05.08.539839

**Authors:** Saman Majeed, Lan Dang, Md Majharul Islam, Olamide Ishola, Peter P. Borbat, Steven J. Ludtke, Elka R. Georgieva

**Author notes:** Correspondence should be addressed to: Steven J. Ludtke or Elka R. Georgieva. These authors contributed equally to this work (Saman Majeed, Lan Dang).

## Abstract

We report our findings on the assembly of the HIV-1 protein Vpu into soluble oligomers. Vpu is a key to HIV-1 protein. It has been considered exclusively a single-pass membrane protein. However, we revealed that this protein forms stable oligomers in aqueous solution, which is an interesting and rather unique observation, as the number of proteins transitioning between soluble and membrane embedded states is limited. Therefore, we undertook a study to characterize these oligomers by utilizing protein engineering, size exclusion chromatography, cryoEM and electron paramagnetic resonance (EPR) spectroscopy. We found that Vpu oligomerizes via its N-terminal transmembrane domain (TM). CryoEM analyses suggest that the oligomeric state most likely is a hexamer or hexamer-to-heptamer equilibrium. Both cryoEM and EPR suggest that, within the oligomer, the distant C-terminal region of Vpu is highly flexible. To the best of our knowledge, this is the first comprehensive study on soluble Vpu. We propose that these oligomers are stabilized via possibly hydrophobic interactions between Vpu TMs. Our findings contribute valuable information about this protein properties and about protein supramolecular complexes formation. The acquired knowledge could be further used in protein engineering, and could also help to uncover possible physiological function of these Vpu oligomers.

## Introduction

The HIV-1 encoded Vpu protein fulfils important functions aiding virus adaptation and proliferation.^1-6^ Despite its small size of about 80 amino acids (aa), the protein has a multi-domain organization and participates in diverse interactions with cellular components. It consists of a short soluble N-terminus followed by what is believed a transmembrane Helix 1 (TM), and two soluble helices Helix 2 and Helix 3 in the C-terminal domain (Figure 1).^5,7-9^ The current knowledge is that Vpu spans the membrane bilayer through its TM Helix 1; it resides and functions in the plasma membrane (PM) and endomembranes of infected cell; it forms a homo-oligomer (presumably a pentamer) in the membranes of Golgi and intracellular vesicles, but not in the endoplasmic reticulum (ER).^10-13^ Monomeric Vpu associates with human tetherin (BST2 protein) in PM via TM Helix 1 forming a Vpu-tetherin complex, leading to enhancement of tetherin ubiquitination and its trafficking for degradation.^14,15^ Therefore, apparently when associated with cellular membranes, this protein exists in several functional quaternary structures. Recently, we found that in addition to membrane-bound self-association, full-length (FL) Vpu also forms homo-oligomers in aqueous solution outside a lipid/hydrophobic environment.^16^

**Figure 1.**
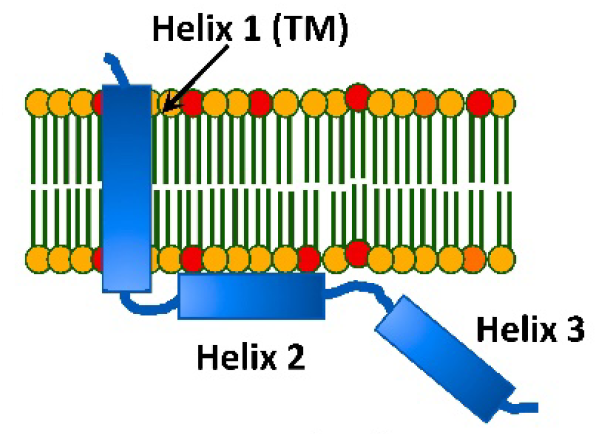
A cartoon representation of the topology of HIV-1 Vpu in lipid membranes: Vpu traverses the membrane via its hydrophobic Helix 1 (TM helix). Helix 2 is membrane-associated; however, other model of Helix 2 not making a contact with the membrane surface was also proposed.

**Figure 2.**
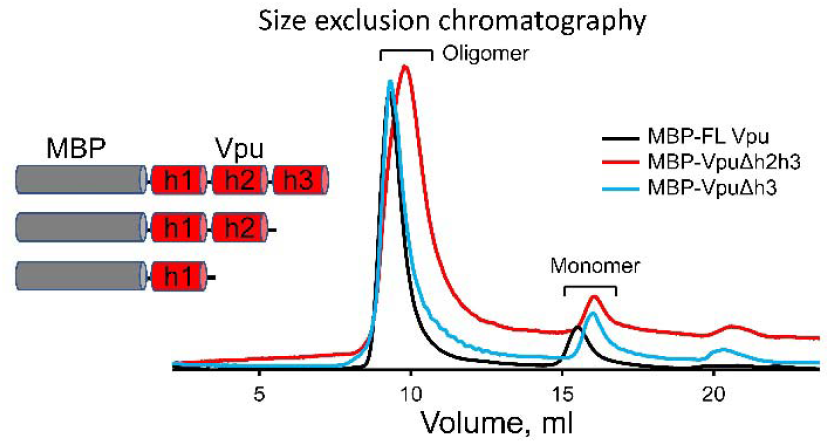
Results from SEC on MBP-Vpu protein in buffered aqueous solution. The SEC results demonstrate that all the MBP-FL Vpu, MBP-VpuΔh3 and MBP-VpuΔh2h3 constructs form oligomers, as they elute predominantly at about 8.5 – 12 ml (depending on the construct) and only small fraction elutes as monomers corresponding to the peak at 16 – 18 ml. The protein concentration was 15 µM for all the three constructs.

Although, such Vpu self-association has been reported before,^10^ we have recently characterized these oligomers in more detail by using a range of biochemical and biophysical techniques, which are analytical size exclusion chromatography (SEC), negative staining electron microscopy (nsEM) and electron paramagnetic resonance (EPR) spectroscopy, revealing that they are rather stable and uniform at protein concentrations of and above 5 µM.^16^ This was an interesting result, given that the prevailing opinion is that Vpu is exclusively a transmembrane protein. Furthermore, a previous study found that soluble Vpu oligomers form even upon the expression of this protein in a rabbit reticulocyte lysate,^13^ which along with our findings brings the question about the possible physiological role of the soluble Vpu form. It could be that the protein is even more functionally versatile than what is currently known about its physiological roles, because of its capacity to transition from soluble to membrane-bound form. Several other unconventional transmembrane proteins from eukaryotic and prokaryotic organisms are known to exist and function in both soluble and membrane-bound states, e.g., the members of Bcl-2 family,^17^ bacterial toxins,^18^ etc., suggesting that this could be the case for Vpu as well.

To advance the understanding of the soluble Vpu oligomers we aimed at characterizing them in greater detail by using the MBP-Vpu construct developed previously^16^ and studying it by cryoEM in combination with SEC and also by continuous wave electron paramagnetic resonance (CW EPR) and double electron-electron resonance (DEER) spectroscopy. The study described here presents a much higher level of sophisticated analyses of the Vpu soluble oligomers, compared to the very coarse analysis we conducted previously.^16^ Here, we provide a more precise model of these oligomers and found that the Vpu protein organizes into most probably hexamers but small fraction of heptamers could also be present, based on cryoEM data. These findings do not contradict our initial assumption of possibly pentameric oligomer organization,^16^ since our previous analyses were based on considerably lower resolution nsEM data. Both cryoEM and EPR results point to significant dynamics and heterogeneity of the distant C-terminus of Vpu, which we reveled for the first time. Thus, the current study not only confirmed with high confidence that in aqueous environment Vpu forms stable oligomers, but also reports for the first time on structural details of these oligomers.

## Materials and Methods

### Protein constructs design, cloning, expression, and purification

In this study, we used three chimeric protein constructs of HIV-1 Vpu fused to maltose binding protein (MBP), which are MBP-Vpu full length (MBP-FL Vpu), MBP-Vpu with truncated Vpu helices 2 and 3 (MBP-Vpu-Δh2h3), and MBP-Vpu with truncated helix 3 (MBP-Vpu-Δh3). The MBP-FL Vpu was the same as one used previously.^16^ The truncated constructs were generated by introducing a stop codon after amino acids R31 and G54, respectively, in the DNA of FL Vpu. The DNAs encoding all MBP-Vpu variants were cloned in the bacterial expression vector (pET15b) at NcoI/BamHI cloning sites. High-throughput prokaryotic expression vectors obtained after cloning i.e. MBP-FL Vpu (MBP-Vpu)/pET15b, MBP-Vpu-Δh2h3/pET15b, MBP-Vpu-Δh3/pET15b were used further for protein expression. For cloning, protein expression, and purification by double-affinity chromatography, *i*.*e*. Nickel (Ni^2+^, Ni)-affinity chromatography and amylose affinity chromatography, the previously developed protocols^16^ were utilized.

The purity of all proteins was assessed by using sodium dodecyl sulfate gel electrophoresis (SDS-PAGE) and western blotting (WB), as described in Majeed *et al*. ^16^

### Assessing the protein oligomeric state using size exclusion chromatography

We used size exclusion chromatography (SEC) to assess the oligomerization state of the purified MBP-FL Vpu, MBP-VpuΔh2h3, and MBP-VpuΔh3 proteins, as described previously. A buffer composed of 50 mM sodium phosphate (NaPi) pH 7.4, 150 mM NaCl, 200 µM TCEP and 10% glycerol was used to equilibrate the column prior to protein sample injection. Then 500 µL of protein sample was injected into the column and eluted by running a SEC program with 0.5 ml/min flow rate. For these experiments, we used a Superdex™ 200 increase 10/300 GL size-exclusion column (*GE Healthcare*) plugged into an AKTÄ explorer 100 (*Amersham Biosciences*) protein purifier system.

### CryoEM grid preparation and data acquisition

Purified MBP-FL Vpu at concentration of 10 µM in a buffer solution containing 25 mM Tris pH 7.4, 150 mM NaCl and 100 µM TCEP was used in cryo-EM experiments. Quantifoil® R 1.2/1.3 Holey Carbon on copper 200 mesh base grids and Quantifoil® R1.2/1.3 micromachined holey carbon on 200 mesh gold grids were used for vitrification of MBP-FL Vpu. Grids were negatively glow discharged at 15uA for 20 seconds using PELCO EasiGlow™. 3.5μl of protein sample was applied to the grids in the humidity-controlled chamber of FEI Mark IV Vitrification Robot. The grids were then blotted on both sides for 1.5 to 2 seconds and plunged into liquid ethane to vitrify.

The MBP-FL Vpu data was collected at the Baylor College of Medicine Cryo-EM ATC using a Thermo Scientific™ Glacios™ Cryo-transmission electron microscope, operated at 200 keV optics. The data was collected with a Falcon IIIEC direct electron detector in counting mode, at a magnification of 120,000, corresponding to a pixel size of 1.200 Å. The dose rate was set to 5.44 e-per Å2 per second. Data was collected using Thermo Scientific™ Smart EPU software. A total of 478 micrographs were collected in one session.

### CryoEM data processing

The raw movie frames were aligned for motion correction in RELION4.0.^19^ Motion-corrected images were then imported to EMAN2^20^ suite for processing. Image processing followed standard EMAN2 single particle reconstruction pipeline^21^, cross-validated using RELION4.0 as outlined briefly below.

Roughly 20,000 putative particles were selected using deep learning-based particle picking in EMAN2.^22^ False positives were manually removed to produce a final set of 14,657 particles used for all subsequent analyses. 128 class averages were produced using the bispectrum-based unsupervised class averaging method in EMAN2. The class averages were then used to generate an initial model by stochastic gradient descent for 3-D refinement. Particles were refined using standard refinement with “gold standard” resolution estimate^23^. 3D refinement was also performed using RELION4.0 for cross validation purposes, with good agreement when using the EMAN2 starting model. Multiple additional refinements were attempted using several different starting models and multi-model refinement methods. Multi-model refinement ^24^ into 4 classes produced two distinct structures. The particles associated with these two structures were then independently refined producing the two maps shown in Figure 3. The newer GMM variability method ^25^ was also applied, and identified the same discrete separation of the data into two groups, along with local variability at the level of roughly the 25 Å resolution achieved in the refinements. The resolution of these maps is not limited by raw image quality, which clearly extended to subnanometer resolution, but rather to the conformational variability of the assemblies within the two identified subsets.

**Figure 3.**
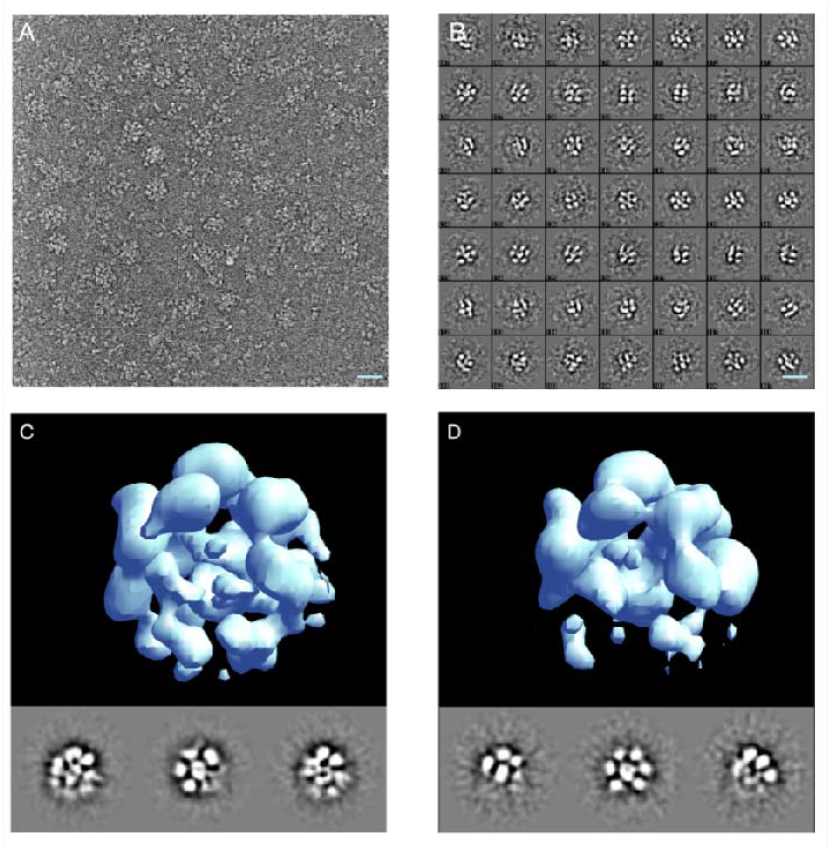
Single particle cryoEM analysis of MBP-FL Vpu soluble oligomers. (A) A micrograph showing representative oligomers and dissociated monomers; (B) representative unsupervised 2D class averages from 14,657 particles; (C-D) single-particle reconstruction of the two identified particle populations with 11,541 (C) and 2815 (D) respectively. The scale bar is 20nm.

### Estimating the number of monomers in the self-associated protein oligomers based on cryoEM structures

Since the major densities observed in the 3-D map are strikingly similar in size to the known MBP structure, the number of monomers in the complex was determined by simply docking into each of the masses of appropriate size in each map. At this resolution structures lack sufficient features to determine subunit orientations with any level of accuracy. Maps are presented as isosurfaces, and significant size variations are possible with different choices for isosurface level. To determine an appropriate isosurface level for docking and visual presentation, the PDB structure of MBP was converted to an electron density map at similar resolution, then the isosurface was selected such that the contained mass agreed with the PDB mass using the standard protein density of 1.35 g/ml (0.81 Da/A^3).^25^ The isosurface of the CryoEM map was then adjusted so the individual docked subunit volumes roughly match the size of the PDB-based monomer structure. An additional transparent isosurface is presented at a lower threshold in Figure 4 to show some of the weaker densities present in the structure representing higher variability regions. As an additional internal control, the volume ratio of the MPB to undocked density in the CryoEM maps is consistent with the relative mass ratio of MBP to Vpu within the margin of error. This increases our confidence that the observed oligomer consists of 6 MPBs. While the larger of the two observed structures arguably has sufficient overall mass to account for 7 MBP/Vpus there isn’t an ideal docking location for a seventh MBP in this structure. Much of the additional mass seems disordered.

**Figure 4.**
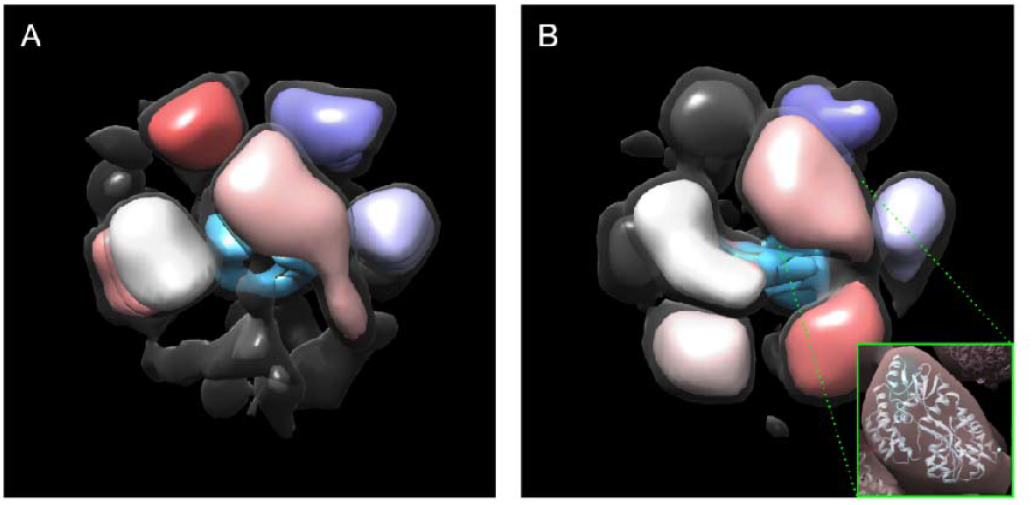
Modeling the oligomeric state of MBP-FL Vpu. (A-B) Maps of the two oligomeric states with populations of 80% and 20% are shown in Figure 3 C and D, respectively. The regions where MPB could be docked are colored in pink, gray, shades of purple, and red. The electron density, which most likely represent Vpu oligomerization core is colored in blue. Views are from different angle than those in Figure 3. A lower isosurface threshold presented in transparent gray shows the lower-density connections in the reconstruction. The inset shows an MBP molecule (PDB: 1PEB) docked into the pink density region of the 3D model as a representative example.

### Protein spin labeling and EPR spectroscopy

Two MBP-Vpu single cysteine mutants labeled with MTS spin label as previously^16^ were used in EPR experiments. The mutations Q36C and N55C were located in the C-terminus of Vpu on opposite sides of Helix 2. Amino acid numbering is that for FL-Vpu.

Continuous wave (CW) EPR experiments were done in the Georgieva lab at TTU with an X-band Bruker EMXmicro spectrometer. The EPR spectra were recorded at room temperature using magnetic field modulation amplitude of 1.5 to 2 Gauss and 0.5 mW microwave power.

Pulse dipolar EPR (PDS) experiments used standard four-pulse double electron-electron resonance, DEER,^26-28^ and DQC.^26^ Both were conducted at ACERT (Cornell University) at 60 K using a home-built Ku-band pulse EPR spectrometer operating at MW frequency around 17.3 GHz.^29,30^ DEER experiment setup^31^ used for electron spin-echo detection a π/2-π-π pulse sequence with 32-ns π-pulse widths applied at the low-field edge of the nitroxide spectrum. A 16-ns pump π-pulse was tuned to 70 MHz lower frequency pumping at the central peak of the spectrum. A 32-step phase cycle ^32^ suppressed all unwanted echo contributions to the signal and some other instrumental artifacts, leaving only three unwanted dipolar pathways produced by the 4-pulse DEER sequence.^33^ Nuclear electron spin-echo envelope modulation (ESEEM) caused by the matrix protons was suppressed by summing up the data traces recorded in four sequential measurements where initial pulse separations and the start time of detection were advanced in subsequent measurement by 9.5 ns that is by a quarter period of the 26.2-MHz nuclear Zeeman frequency of protons at 0.615 T corresponding to 17.3 GHz working frequency.^34^

DQC experiments (conducted on N55C) used 6-pulse sequence with 6 ns π-pulses and 64-step phase cycle isolate the dipolar signal. No unwanted dipolar pathways are generated in this sequence. The short distances between the labels produced very fast dipolar evolution appearing as a narrow peak in the center of the symmetrical in time dipolar signal. An apparent small background was due to couplings between labels at longer distances as can better be seen in DEER data. The additive background contribution is most likely very small in this case and can be left in place to be processed together with the main part of the dipolar signal.

Primary echo decays were routinely recorded for all samples. Supporting Figure 1 shows echo decays for both mutants in protonated and deuterated buffers.

Some of time-domain DEER data traces *V*(*t*) were processed to distances. All traces were conditioned using standard approaches^26,31,35,36^ and then reconstructed into distance distributions *P*(*r*)’s by either Tikhonov regularization method or an SVD application accessible via ACERT website. The DEER signal record *V*(*t*) was corrected for the decay caused by intermolecular dipole–dipole couplings by fitting the latter points (about half of the record) of ln[*V(t*)] to the straight line extrapolated to zero time and subtracted from ln[*V(t*)], giving the antilog *u*(*t*) = *d(t*) + 1, where *d*(*t*) is contributed by the dipolar evolution of DEER signal. Then, *u*(*t*) normalized as *u*(*t*)/*u(*0) gives typical presentation of DEER data, (cf. Figure 7). Another data form, *v*(*t*) = *d(t*)/*u*(0) gives background-free “dipolar” data used as an input in reconstructing distances between the coupled electron spins. The DEER data for the two spin-labeled residues are shown in Supporting Figures 2 and 3.

## Results

### The Vpu soluble oligomers are a result of Helix 1 (TM) self-association

To establish with certainty which region of Vpu contributes to its oligomerization in solution, in addition to MBP-FL Vpu, we produced two truncated versions of Vpu that are MBP-VpuΔh3 and MBP-VpuΔh2h3 (with deletions of either Helix 3 or both Helices 2 and 3). We studied these protein versions with SEC and found that all MBP-Vpu variants elute as oligomers of close molecular weight (Figure 2). MBP-Vpu oligomers were formed even when both Helices 2 and 3 were deleted, although the corresponding elution peak at 8.5 – 12 ml was broader than the elution peaks of MBP-FL Vpu and MBP-VpuΔh3. The estimated full width at half maximum (FWHM) for these peaks were 0.74, 0.85 and 1.5 (in ml), respectively, for the FL Vpu, VpuΔh3, and VpuΔh2h3. The observed elution peak broadening for Vpu lacking both Helix 2 and Helix 3 possibly indicates that just Helix 1 is insufficient for stabilizing the Vpu oligomers, leading to increased heterogeneity in the oligomeric order. However, increased hydrophobic interacttions between Vpu and column beads due to more exposed Vpu Helix 1 cannot be ruled out as areson for the broadening of the SEC peak. Still, no complete oligomer disassembly occurs, as the intensity of the SEC elution peak at 16 – 18 ml remains unchanged (Figure 2). On the other hand, a little if any difference was observed in the SEC profiles of MBP-FL Vpu and MBP-VpuΔh3, suggesting the essential role of protein region comprising Helice 1 and likely Helix 2 in Vpu self-association and oligomer stabilization, which is in agreement with what was proposed for Vpu oligomer in lipid environment.^12^

As we reported previously, the assessed by SEC MBP-Vpu oligomers have a molecular weight greater than 200 kDa and elute well after the void volume for the SEC column we used (Supporting Figure 1).^16^ We furter found that the MBP moiety does not contribute to Vpu oligomerization, as when we fused MBP to another single-span viral membrane protein, namely the coronavirus E protein, no oligomerization was observed (Supporting Figure 2).

### CryoEM provides a solid evidence about the formation of stable soluble MBP-FL Vpu oligomers

In a previous study, we utilized nsEM to observe stable soluble oligomers of MBP-Vpu.^16^ These oligomers were typically formed at protein concentrations exceeding 5 µM, which indicated that cooperative mechanisms may have contributed to this process. MBP-FL Vpu oligomers were imaged using cryoEM to confirm that the well-defined soluble oligomers observed by nsEM are reproducible under cryoEM conditions that utilize fast freezing of the sample. The cryoEM freezing process occurs on the timescale, which is too fast for water ice to form and grow and for kinetically slow processes, such as oligomer disassembly, to materialize,^37^ thereby preserving the native protein state.

To perform structural analysis by single-particle cryoEM, purified MBP-FL Vpu protein was used. The oligomerization of MBP-FL Vpu in buffer solution was confirmed by SEC (Figure 2) prior to cryoEM imaging. CryoEM data collected in the present study had sufficient contrast to enable clear visualization of oligomeric proteins in the raw micrographs (Figure 3A). We collected a data set of 14,657 putative oligomeric particle images, sufficient for structural analysis. A representative micrograph containing MBP-Vpu oligomers, and reference-free 2D class averages are shown in Figures 3A and 3B, respectively. The micrograph exhibits mostly oligomeric assemblies similar to those previously observed in nsEM,^16^ but a small fraction of lower order oligomers and possibly monomers were also present. Notably, the majority of the self-associated protein particles were of a visually similar size, and only small fraction of non-specific larger aggregates could be found. When further analyzed, the visibly uniform particles readily group into unsupervised classes of a similar size implying a self-consistent quaternary structure of size-limited protein oligomers.

To confirm whether the class-averages represent different views of a single 3-D quaternary structure or represent multiple states, we conducted de-novo reconstructions using EMAN2,^19^ performing several cross-validations and classification attempts. The analysis produced two self-consistent 3-D structures at low resolution (∼25 Å), representing different populations of the observed particles. Out of a total of 14,657 particles, about 80% were associated with one larger-size structure (shown in Figure 3C) with several low-intensity features, whereas the remaining about 20% of the particles correspond to a similar but slightly smaller-size structure (Figure 3D). The densities populating the centers of the structures in Figures 3C and 3D, presumably correspond to the Vpu TM oligomer core. The remote low densities observed outside of the structures, more visible in Figure 3D could possibly be attributed to a particular arrangement of the distant Vpu C-terminal helices. We made attempts to further classify the particles or characterize the remaining variability in the system not accounted for in the two classes by using a deep-learning Gaussian Mixture Model (GMM) approach.^25^ However, this effort did not yield useful insights beyond confirming the overall positioning of the major densities in both structures with the persisting variability at the level of the ∼25 Å resolution. The high monomer flexibility due to MBP-Vpu linker and currently unknown Vpu folding state in the soluble form, and variations in the oligomeric order majorly contribute to the low structural resolution.

The obtained structural models of MBP-Vpu do not provide sufficient structural resolution for further accurate docking of the Vpu oligomeric core; at this resolution and with limited constraints on the isosurface threshold level, it can only be generally placed within the total structure density. On the other hand, due to larger size and greater stability, the densities of MBP in the 3-D structures, could be modeled through docking to estimate the number (N) of MBP moieties, hence the number of MBP-Vpu protomers. We were able to dock the MBP moieties into six MBP-sized domains in both CryoEM maps. The docked MBP moieties in Figure 4 are colored, whereas the remaining densities, shown in blue should account for the Vpu oligomerization core. The remaining dark gray density, which was not attributed to neither MBP or Vpu oligomerization core, is more prominent in the structure shown in Figure 4A, Therefore, it may well be that this structure can accommodate a seventh MBP-Vpu protomer, which might imply the existence of heptamers as well. Then, heptamer-to-hexamer oligomer equilibrium cannot be ruled out.

### Protein spin labeling and EPR spectroscopy confirm that the distant C-terminus of Vpu is highly dynamic

Before undertaking full-fledged multi-site spin labeling EPR experiments to map protein dynamics and solve spatial organization of monomers in the Vpu oligomer, we limited this task to using two carefully selected spin-labeled sites in an effort-saving approach. We reused the generated previously single cysteine mutant Q36C^16^ in the linker at the beginning of Helix 2 N-terminal side and added new labeling site N55C in the linker connecting Helix 2 to Helix 3 (Figure 5A). The selected for EPR studies residues are located away from the MBP moiety of MBP-Vpu, thus we believe they are not affected by the presence of MBP. Therefore, we studied these residues in the context of FL MBP-Vpu for consistency with the cryoEM data and data analyses.

**Figure 5.**
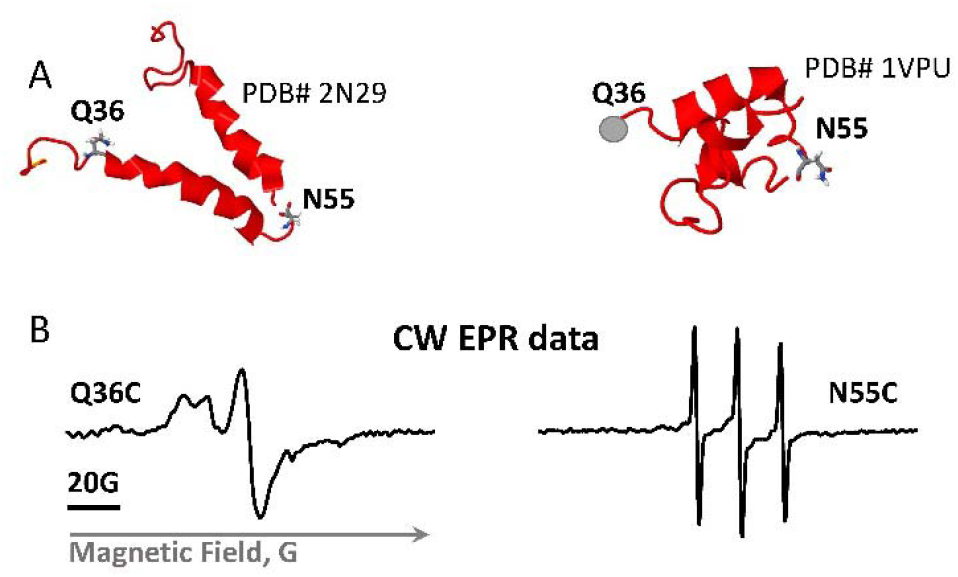
Vpu spin labeling and CW EPR results. (A) Shown are two NMR structures of the soluble C-terminal domain of the Vpu protein with the residues mutated to cysteines and spin-labeled. The residue Q36C is located at the beginning of amphipathic Helix 2, residue N55C is in highly dynamic loop connecting Helices 2 and 3. (B) EPR spectra of spin-labeled residues Q36C and N55C show very different mobility of spin labels at these sites.

The broad CW EPR spectrum recorded for spin-labeled Q36C mutant (Figure 5B) reinforced the previous conclusion that this protein region has restricted motional dynamics.^16^ This is in line with the SEC results showing that the linker connecting Helices 1 and 2 play a role in Vpu soluble oligomer stabilization. This may require specific structural arrangement of the protein region containing residue 36. Could be that this linker is ordered, possibly in contact with the oligomer-forming Helices 1. On the other hand, the CW EPR spectrum for the spin-labeled residue N55C has narrow lines (Figure 5B), indicating that this protein region is highly mobile and possibly unstructured,^38-42^ in agreement with the structures of monomeric Vpu C-terminal (Figure 5A).^43,44^ However, a minor population of protein conformation with restricted motional dynamics at this site contributing broad low-intensity lines to the EPR spectrum of this spin-labeled residue while cannot be ruled out was not immediately apparent in the signal dominated by the high-intensity lines of the predominant conformation.

The pulse EPR (DEER) data for the Q36C mutants were already recorder and published.^16^ It was of more interest to conduct DEER distance measurement for the newly spin-labeled residue N55C, probing conformations of this protein region. For the DEER measurements of this mutant, we used partial or full solvent deuteration by substituting H_2_O for D_2_O and glycerol for glycerol-*d*_8_ in the buffer aiming to increase the signals and extend the upper range of measurable distances at least for more water exposed spin-labeled sites due to increased spin label phase memory relaxation time, *T*_m_.^45-47^

Unlike Q36C,^16^ the residue N55C is favorable to spin labeling. Long *T*_m_‘s and larger primary echo amplitude compared to Q36C (Supporting Information) show that this residue is more accessible to aqueous phase. The two recorded DEER traces for spin-labeled mutant N55C and evolution times of 1 µs and 3 µs (Figure 6A) revealed very different form-factors of signal decays. The clearly bimodal character of DEER signal trace for the shorter evolution time (*t*_m_) and the respective distance distribution are shown in Figures 6B and 6C. For the 3 µs data, however, one can see what appears mostly as a slow “uniform” background with a small narrow feature in the beginning of the record. To produce such signal in a spin pair widely distributed distance above 80 Å would be necessary.^48^

**Figure 6.**
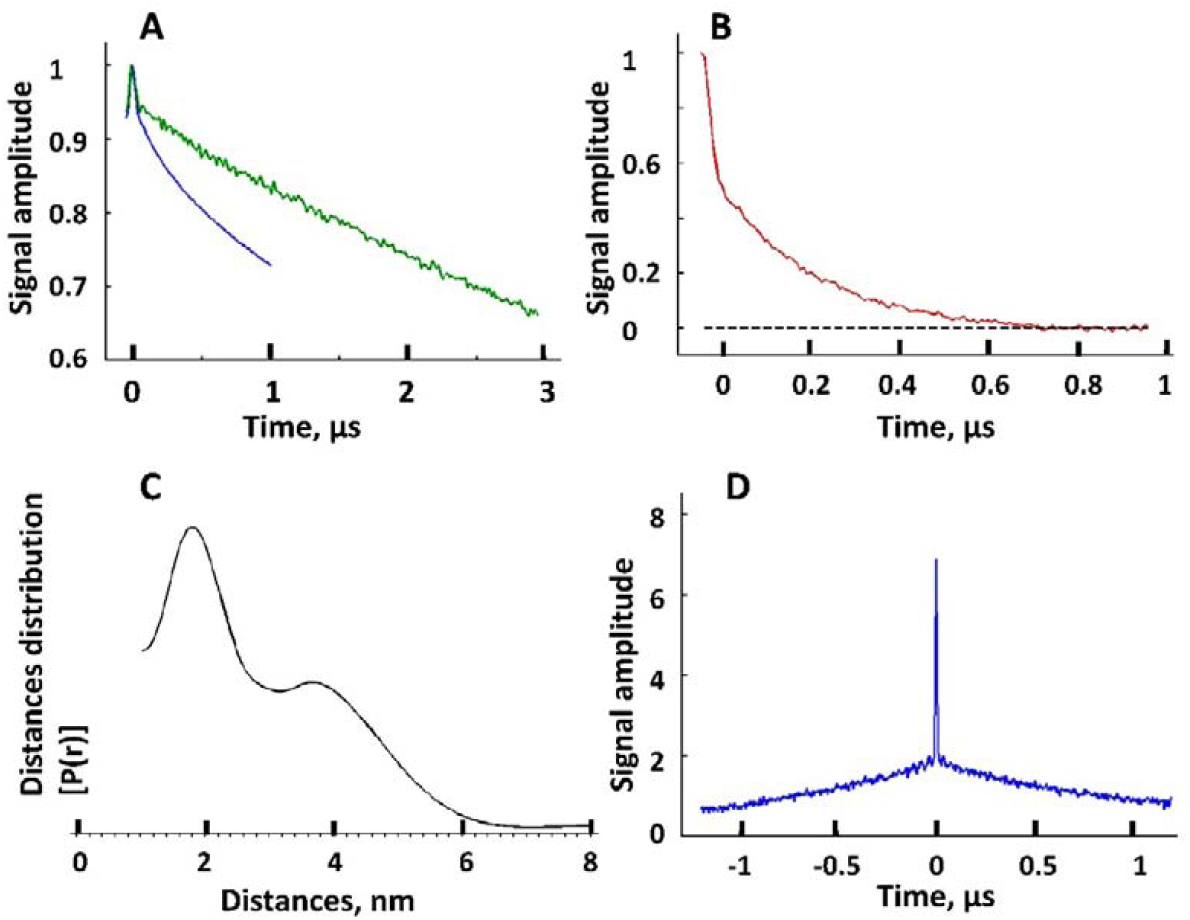
The DEER data and reconstructed inter-spin distances for Vpu spin-labeled at residue N55C. (A) Raw DEER data recorded for two different evolution times, 1 µs (blue) and 3 µs (green). (B) Background subtracted DEER signal for 1 µs evolution time data in A. The black dashed line indicates the zero-signal amplitude. (C) The reconstructed inter-spin distances for the data in B. (D) The intense narrow peak in DQC data originates from very short distances on the order of 10 Å for some spin-labels at N55C in Vpu oligomer. These distances are too short to be sensed by DEER whose sensitivity rolls off quickly below 20 Å.

**Figure 7.**
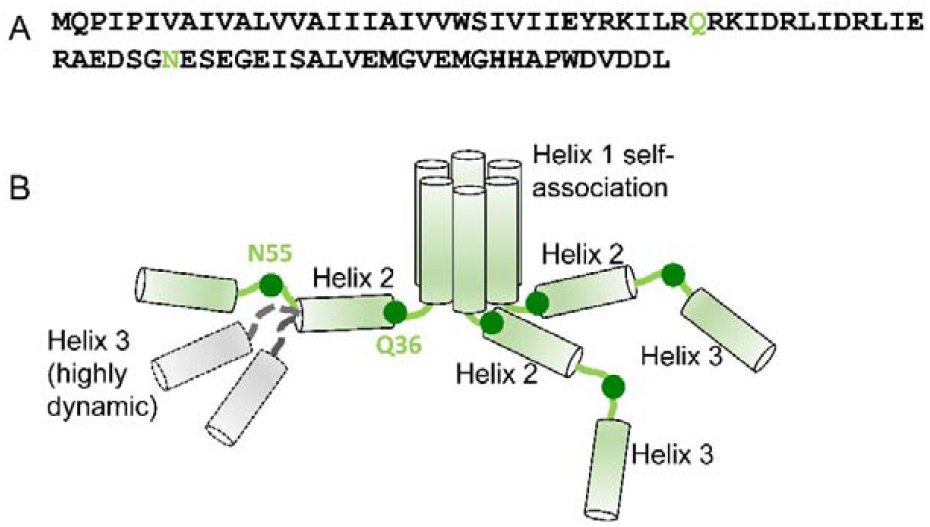
Proposed model of the soluble Vpu oligomers. (A) Amino acid sequence of FL Vpu. The linker between Helix 1 and Helix 2 is highlighted in gray, and the residue Q36 is in green. (B) Hexametric arrangement of Vpu monomers: Helices 1 from six protomers come together to form the oligomer core. The linker between Helix 1 and Helix 2, and Helix 2 in each protomer are arranged in a way to stabilize the oligomer. The residues Q36 and N55 are shown as dark green circles. The residue Q36 is possibly oriented toward the oligomer core and occluded. Helix 3 is widely spread and conformationally heterogeneous. Helices 2 and 3 for only three protomers are shown for clarity. Helix 3 is highly dynamic and heterogeneous in conformation—Possible alternative conformations of this Vpu region are shown for just one protomer for the sake of clarity. Also, for the clarity of representation, the model shown here does not account for the possible interactions between Helices 3 from neighboring Vpu protomers.

These observations suggest that spin labels in close proximity or at moderate distances in Figure 6C and Figure 6D have shorter relaxation times than a fraction of labeled sites, which may be located far away from these apparent spin clusters by the nature of the oligomeric assembly. They could be residing on less restricted “floppy” domains, so that only small dipolar evolution occurs on a limited time-scale. For long evolution times these slowly-relaxing spins-labels contribute most of the echo signal and long distances due to these sites dominate the DEER signal. It is unlikely that these slow decays are caused by intermolecular spin interactions since oligomer spin clusters are confined to widely separated complexes. In such cases analogues to spin-labeled micelles, intermolecular contributions to DEER signal are small.^48^

This result suggests that the Vpu region containing the loop between Helices 2 and 3, and possibly Helix 3, are sampling multiple conformations outside the Vpu oligomer core. This is in line with cryoEM results (Figures 3 and 4) indicating that this protein region is not well resolved in the structures.

However, in addition to this broadly distributed distances, for the residue N55C we also observed a measurable contribution of very short inter-spin distances at or below 10 Å reported by double quantum coherence (DQC),^49^ as shown in Figure 6D. Apparently, these very short distances correspond to a not yet quantified fraction of labels occupying sites locked in distinct conformations. They possibly contribute broad and difficult to quantify lines in the EPR spectrum of residue N55C in Figure 5B. Based on these results, we may conclude that some of Helices 2 and possibly Helices 3 from different protomers interact with each other and/or the oligomeric core, bringing into proximity the residues N55 in the Vpu oligomer. This could explain the presence of peripheral low electron densities in the cryo-EM structures seen in Figure 3D.

## Discussion

Vpu is a HIV-1 encoded accessory protein, which has been linked to several processes important for the virus functions in the infected cells.^4,6,50,51^ However, at present, the Vpu structures underlying these functions are not characterized in sufficient detail. In addition, in cellular membranes, Vpu is thought to function in both oligomeric and monomeric states,^10^ but the factors governing the quaternary structure variations have not been adequately identified and studied. Furthermore, we^16^ and others^13^ found that this protein forms soluble oligomers outside of membrane or hydrophobic environment. This is atypical behavior for a transmembrane protein, although other such proteins alternating between soluble and membrane-bound functional states are known.^17,18^ It may well be that the observed soluble oligomers of Vpu are purely a result of fortuitous physicochemical properties, as currently there is insufficient evidence to confirm or rule out their physiological relevance. One plausible role could be that soluble oligomers serve a shuttle of this protein between cellular compartments. Nevertheless, the soluble Vpu oligomers present an interesting supramolecular construct to study interaction and folding of certain polypeptides, cooperativity, and formation of protein superstructures. This motivated us to seek out understanding of their structural properties and the mechanism of assembly. To this end, we utilized a combination of several experimental techniques that are cryoEM, EPR, SEC and protein engineering to maximize the acquired information about the soluble Vpu oligomers. Given that FL Vpu monomer have molecular weight of ca. 9 kDa, to maximize the protein size-sensitive cryoEM studies, we benefited from the MBP-Vpu construct having an increased effective molecular weight and size. This approach has been used widely in cryoEM studies on small proteins.^52,53^ The obtained cryoEM data were of adequate quality to build a 3-D structural model of MBP-Vpu oligomers and the best fit of this model was obtained for MBP-FL Vpu hexamers, but heptamers cannot be ruled out. We also found that the oligomer formation is a result of Vpu TM self-association, and Helix 2 has stabilizing effect, as observed by SEC. It is not yet known how the regions of TMs are structured in the Vpu oligomer. The CW EPR and DEER data based on the singly spin-labeled residue N55C in Vpu C-terminal domain indicated that this Vpu region is highly dynamic, sampling a wide range of conformations, and the labeled sites at the C-termini of protein protomers are mostly widely separated.

Thus, the results from this and previous studies provided unambiguous evidence that Vpu forms well-defined and stable oligomers in solution. Further studies, will make available more information on the structure of these oligomers and could possibly serve as a ground to uncover a physiological function linked to them.

Based on the results from this study, we propose a model of Vpu soluble oligomers in which the regions of Helices 1 self-associate to form a hydrophobic core of possibly a hexameric quaternary structure, which is further stabilized by the favorable arrangement of amphipathic Helices 2 and linkers between Helix 1 and Helix 2 of each protomer (Figure 7). The linker between Helices 1 and 2 is possibly occluded, which could explain the difficulties to spin label the residue Q36C, as well as the obtained slow-motional CW EPR and DEER data for this residue.^16^ Helices 3 are predominantly spread apart and are highly dynamic.

It is also interesting to mention that a previous study found that the phosphorylation of Ser 52 and Ser 56 in Vpu polypeptide enhances the protein oligomerization in detergent.^54^ However, our results in solution suggest that such phosphorylation is not needed, as under our experimental conditions the equilibrium was shifted exclusively toward stable FL Vpu oligomers and the oligomerization was unaffected when the region containing Ser 52 and 56 was truncated (Figure 2). The observed differences might be a result of an altered Vpu conformation in detergent vs. solution.

Furthermore, since Vpu have been considered exclusively a transmembrane protein, several NMR studies report on its monomer structure and oligomerization in lipid environment and a pentamer was proposed as the most probable quaternary organization.^55-57^ In addition, we observed Vpu oligomer restructuring upon transition from soluble to lipid-bound state.^16^ Therefore, it could be that the oligomeric organizations of Vpu in lipid and in solution are not identical, taking into account that the interactions between Vpu hydrophobic Helices 1 in aqueous solution and in lipid environment participate in different interactions.^58^

## Acknowledgements

We greatfuly acknowlege Isaac Forrester for collecting the cryo-EM data used in this study. Oluwatosin Adetyui is acknoledget for participation in intial production of MBP-FL Vpu protein. The study was supported by start-up funds from the Department of Chemistry and Biochemistry at TTU (to ERG), NIH grant R01GM080139 to (SJL, Baylor College of Medicine), and NIH grant 1R24GM146107 to National Biomedical Resource for Advanced ESR Techniques.(ACERT) at Cornell University.

## Author contribution

SM: experiment, data analysis and interpretation, figures, writing the manuscript; LD: data analysis and interpretation, figures, writing the manuscript; MMI: experiment, data analysis; OI: experiment; PPB: experiment, data analysis and interpretation; figures, writing the manuscript; SJL: supervision, funds acquisition, data interpretation; ERG: conception, design, experiment, data analysis and interpretation, figures, writing the manuscript, supervision, funds acquisition. All authors contributed to and agreed with the final version of the manuscript.

## Data availability

The cryoEM datasets generated and/or analysed during the current study are not publicly available as they are too large to be placed in the supplement but are available from the corresponding authors Steven J. Ludtke and Elka R. Georgieva on reasonable request.

## Notes

### Competing Interest Statement

The authors have declared no competing interest.

### Summary of Updates

Introduction and Discussion updated to better clarify the significance of this work. New references provided to support the significance of the new findings and compare this work with prior published works. Supporting Information updated.

