## Supplementary Information for "HIV-1 Vpu protein forms stable oligomers in aqueous solution via its transmembrane domain self-association"

**SEC on MBP-Vpu constructs**

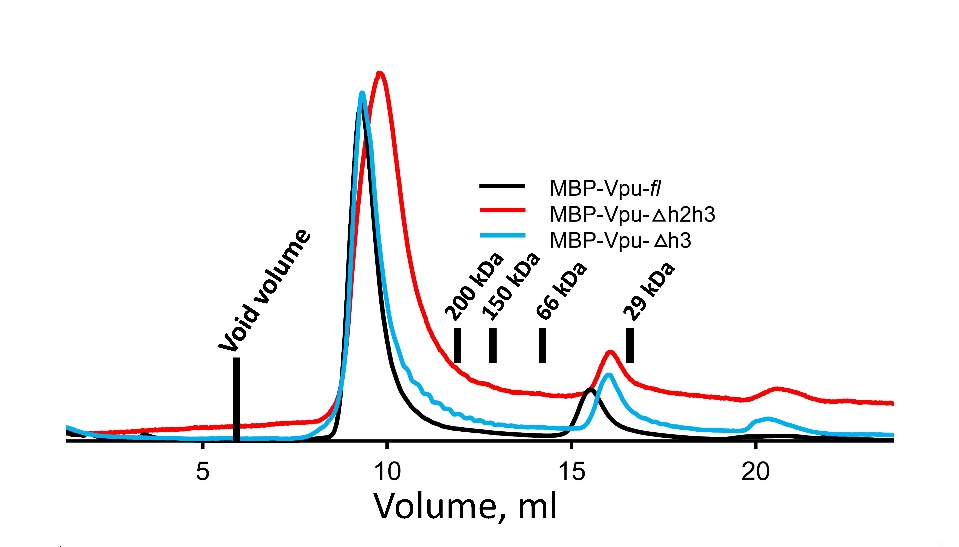

**Supporting Figure 1.** SEC data for the used in this study MBP-Vpu FL and truncated versions. The positions of the elution peaks of protein markers[1] are shown by black bars and the corresponding protein molecular weights. The void volume is typically 20–25% of the column bed volume, which is less than 6 ml (shown in the figure) for the 24 ml column we used.

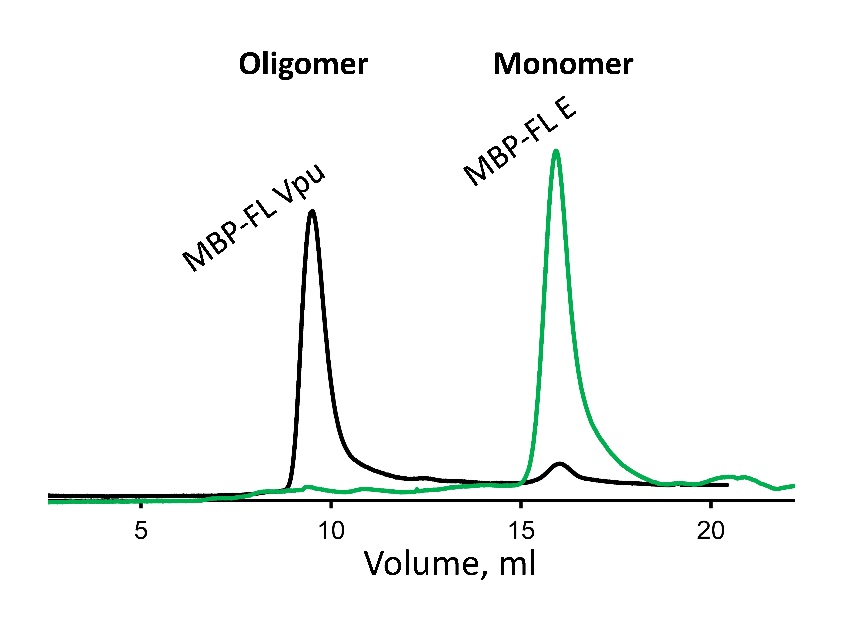

**Supporting Figure 2.** SEC elution profiles for MBP-FL Vpu and MBP-FL E proteins. The experiments for both proteins were conducted under identical conditions. Similarly to Vpu, we used the coronavirus E protein (also a single-pass transmembrane protein) to generate the MBP-FL E chimera construct. This protein was expressed in soluble form and purified using the protocol for MBP-FL Vpu.

**Pulse EPR experiments**

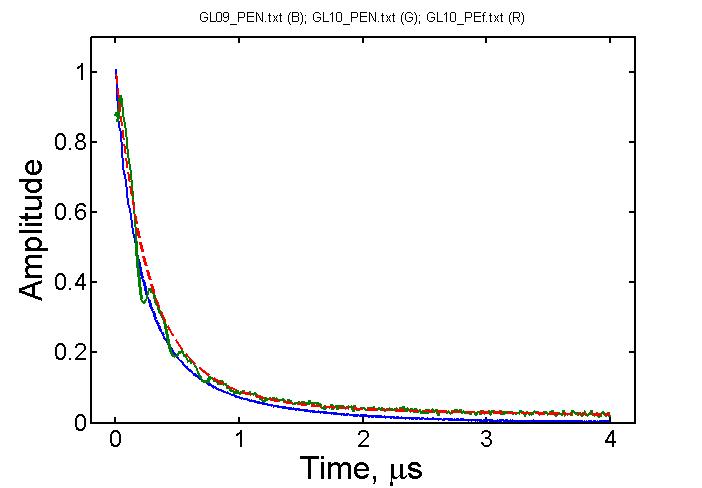

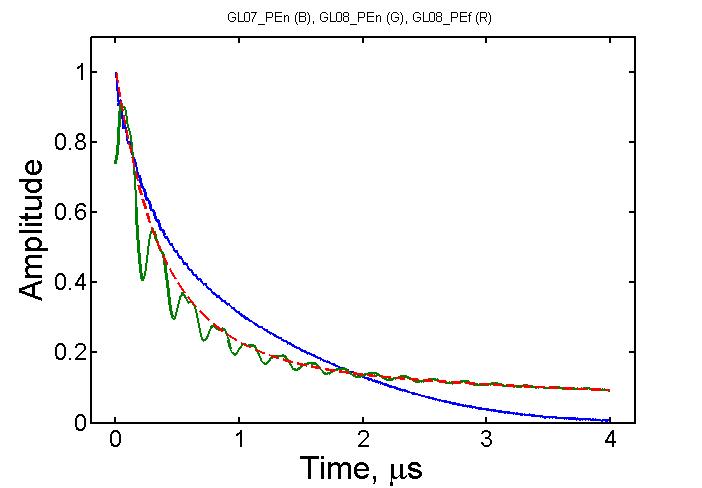

**Supporting Figure 3.** Primary echo decays for Q36C (left) and N55C taken in protonated (in blue) and deuterated solvents (in green) with the latter showing ESEEM from matrix deuterium nuclei. Dashed lines in red approximate the decays for deuterated samples.

There is only small difference in decay rates and deuterium ESEEM is small for Q36C reinforcing the notion that the site is buried. The traces show the presence of spin-labels with longer phase relaxation times not decaying completely to 4 µs. While these could be unremoved free spin label, more likely it is a slow-relaxing fraction of spin label in oligomer folding such as to place some of the label in different types of sites, some possible separated by a long distance. This has implication on DEER traces recorded using longer evolution times *t*_m_. The spin-labels clustered at short and moderate distances and partly buried relax faster, so their dipolar signal could only be well expressed at shorter evolution times. At long evolution times the data are dominated by either free label background or the dipolar evolution of slow relaxing peripheral labels that are more exposed to aqueous medium and separated by long distance from the labels at oligomer core, which contribute the main part of visible evolution of dipolar signals.

A	B

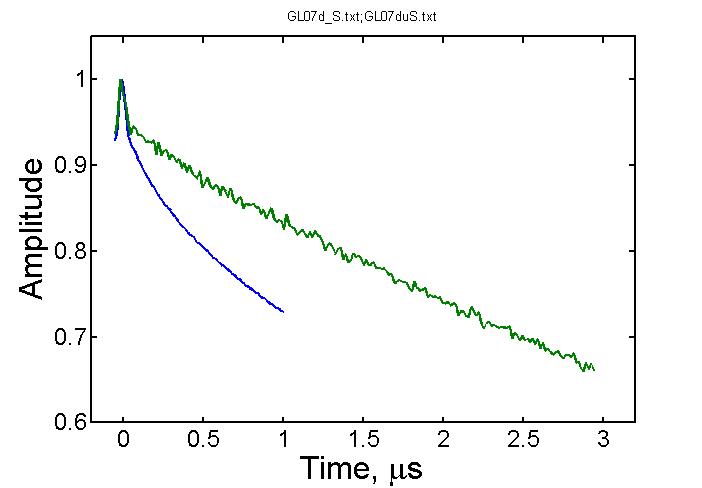

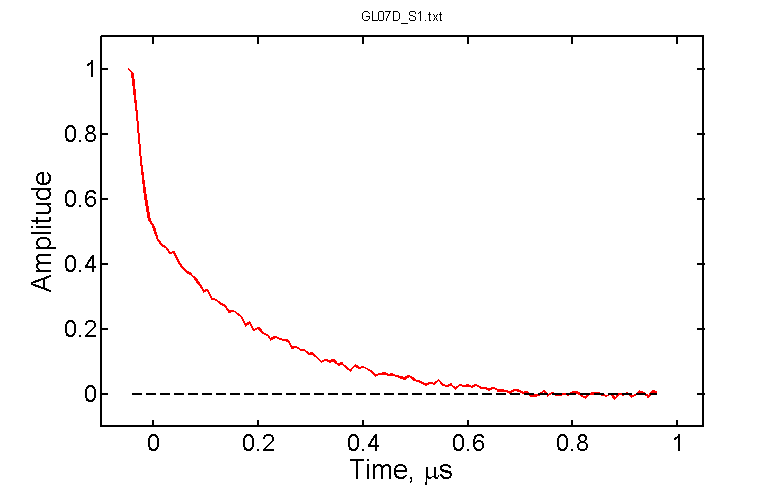

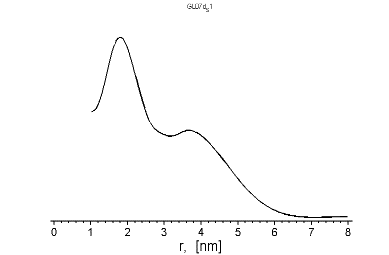

C	D

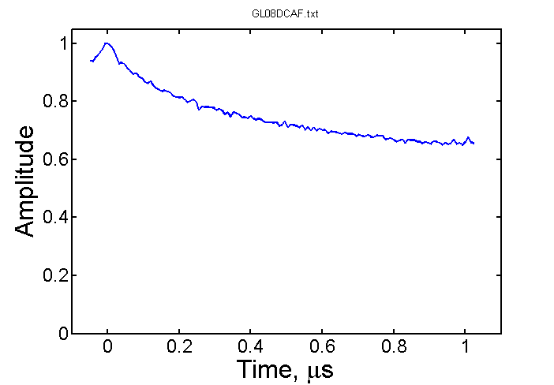

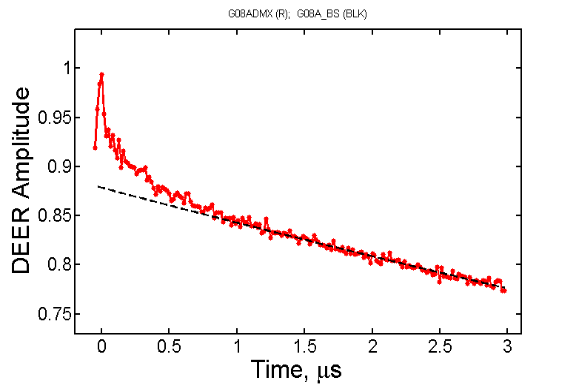

**Supporting Figure 4.** (A) DEER data for N55C in H_2_O buffer at two different evolution times (*t_m_*-s) that are 1 µs (blue) and 3 µs (green); (B) The background-subtracted 1 µs *t*_m_ data from panel A (in red) show complex decaying signal. The dashed black line indicated the zero-signal intensity. The insert in black reveals the bimodal distance distribution reconstructed by Tikhonov regularization; (C, D). The data for different evolution times (at 1 µs *t_m_* in C in blue, and at 3 µs *t_m_* in D in red) for this site in D_2_O buffer. The black dashed line in D correspond to the background signal.

A	B

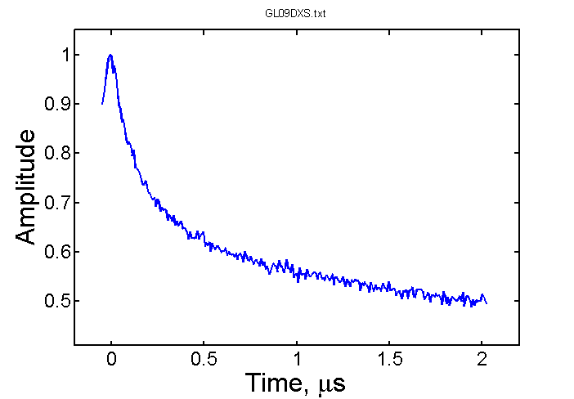

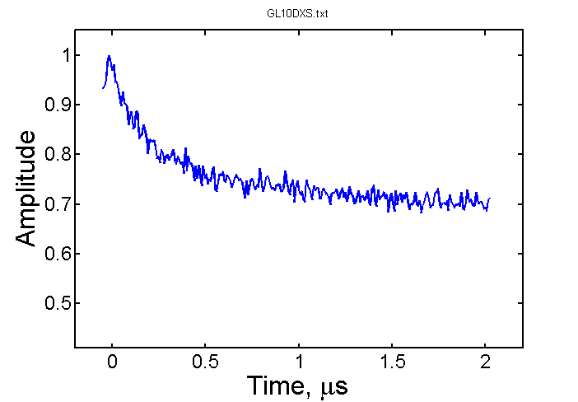

C	D

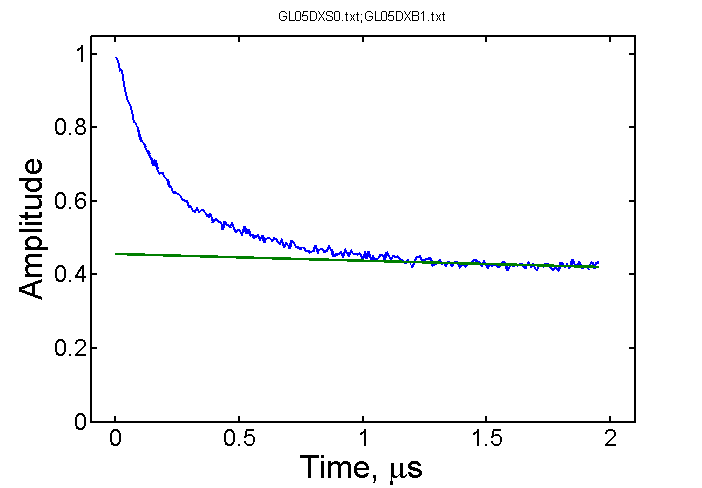

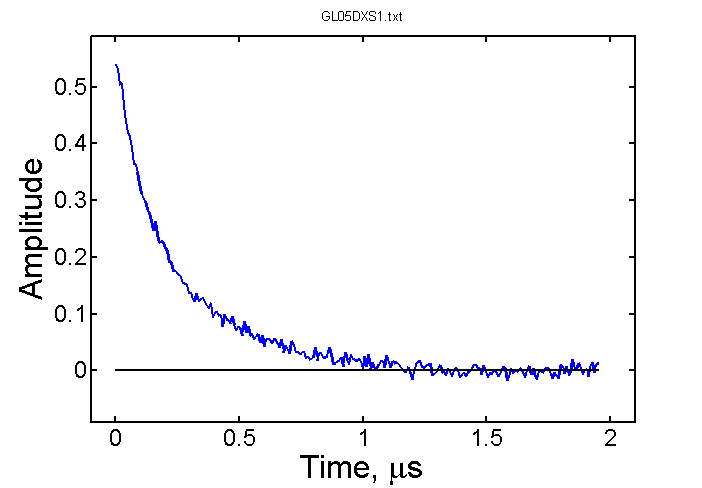

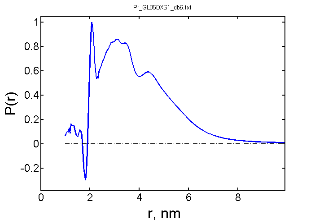

**Supporting Figure 5.** DEER data for Vpu labeled at Q36C recorded in H_2_O buffer (A) and D_2_O buffer (B) are similar but differ in the amplitude of background. The raw DEER data in (C) for another sample in partly D_2_O buffer are shown in (D) with the background subtracted and the remainder processed to distances shown in the insert. The distances reconstruction used SVD in ACERT online software. Very narrow out-of-phase peak originates from a low-level deuterium ESEEM present in the signal. The deuterated sample has similar contributions to distances as the protonated sample. The signal is contributed by more than two spins.

Linear background was subtracted from the DEER signal for residue Q36C. The DEER signal for residue N55C is complex, possibly contributed by short distances for some residues, roughly estimated to be in the range of ~10 Å, as well as by longer distances distributed over broad range due to heterogeneous nature of the labeling site in the linker producing not so well-defined decaying signal, very long distances outside 8 nm could also be present which would not be a total surprise for an oligomer containing several spin labels.

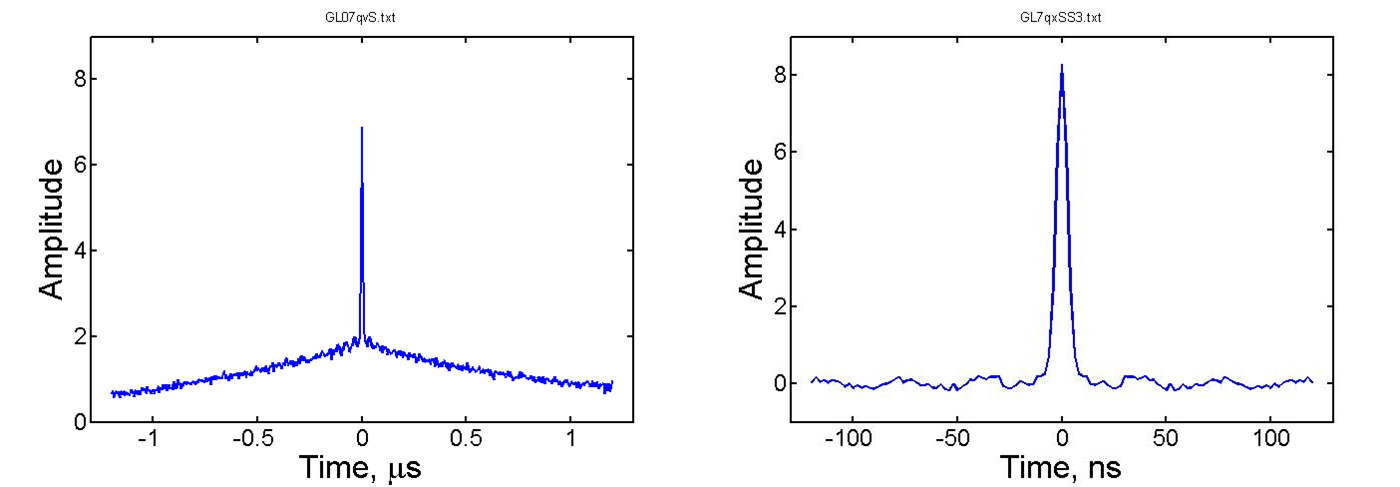

**Supporting Figure 6.** (left) DQC data for N55C in H_2_O. The same data are shown (right) on extended time scale with the slow-decaying underlying signal removed. The fast signal in the center decays in a few nanoseconds. Such fast decay can only be produced by dipolar coupling tens of gauss strong corresponding to distances of 10 Å or shorter.

1.	Majeed, S., et al., *Insights into the oligomeric structure of the HIV-1 Vpu protein.* Journal of Structural Biology, 2023. **215**(1): p. 107943.
